# Consolidation of Sequential Planning

**DOI:** 10.1101/2024.11.01.621475

**Authors:** Oliver Vikbladh, Evan Russek, Neil Burgess

## Abstract

Thriving in changing environments requires the capacity to evaluate novel courses of action^1^. This ability is hypothesized to depend on sequential planning via step-by-step simulations of the future ^2–4^, using cognitive maps or schemas of task contingencies^5,6^. However, it is still unclear if, how and where in the brain such flexible planning is enacted. In parallel, it is thought that consolidation transforms memory representations over time to promote adaptive behaviour^7^. Here, we hypothesize that consolidation strengthens cognitive maps of task contingencies used for simulation during sequential planning. To test this, we developed a novel behavioural task and a new multivariate method for analysis of magnetoencephalography (MEG) data. Using choice and reaction time data we dissociated flexible sequential planning from alternative non-sequential strategies, and identified this behaviour with robust neural markers of step-by-step simulation, localized to the anterior medial temporal lobe. Retesting a week later we showed that consolidation enhanced sequential planning and strengthened markers of sequential simulation in the prefrontal cortex, consistent with systems consolidation theory^8–15^. By revealing that consolidation improves future simulations for flexible planning we open up a new frontier for the investigation of the functional interactions between memory and decision-making.

## Main

How does the brain leverage knowledge about the world to flexibly evaluate actions in novel situations^1^? Recent theories have looked to algorithms from model-based (MB) reinforcement learning (RL) in which flexible evaluation involves sequential planning via step-by-step simulations through cognitive maps of task structure, also known as a ‘’roll-outs’ ^2–4^. It is commonly suggested that the site of these rollouts could be the hippocampus^4,16^, which represents relational cognitive maps ^5,6^, displays non-local sequential activity ^17–19^ and supports flexible sequential evaluation^20,21^. However, definitive evidence for flexible planning through online (i.e. at choice time) rollouts is lacking. Furthermore the hippocampal role in planning appears to be transient^22^. In parallel, theories of memory consolidation propose that memories initially encoded in the hippocampus allow formation of increasingly schematized representations in the prefrontal cortex (PFC)^8–15^. Consolidation is thought to promote adaptive behaviour^7^ but the mechanism by which this occurs is unclear.

Here we address these multiple gaps to test the hypothesis that consolidation transforms and strengthens map-like representations of task contingencies to facilitate more flexible planning via rollouts. With a novel behavioural task and a new multivariate MEG method we show that sequential planning reflects step-by-step simulation at choice, localized to the hippocampus. We then demonstrate that consolidation improves sequential planning and strengthens neural markers of rollouts, particularly in the PFC. Our results suggest that the consolidated representations are relational schemas of task structure, indicating how one task state leads to the next. After learning about changes in the environment, like new obstacles or rewards ^1,23,24^, these representations are used to flexibly adapt decisions by more effectively simulating future outcomes ^4,25^.

### Behavioural Task

We aimed to elucidate the effect of consolidation on flexible planning. However, previous neuroimaging experiments investigating flexible planning ^26–29^ have used flexibility probes which are not specific to sequential planning via rollouts per se. Rather, these probes can also be solved by less flexible strategies, like the ‘successor representation’ (SR), which rely on temporally abstract multi-step representations in place of sequential simulation ^30–32^. While the former entails a sequential unfolding (e.g. State A → State → B State C), the latter operates on compressed representations of subsequent states, without their relative ordering (e.g. State A → States B&C). For estimating the expected future reward in a State A, both strategies can integrate new information about which states are rewarded (e.g. State C now contains reward), known as Reward Revaluation. However, only the sequential strategy is flexible enough to make use of new information about the transition structure (e.g. there is now an obstacle between State B and State C, so the reward is unreachable), known as Transition Revaluation ^31,32^. We therefore created a new task to measure sequential planning, with high power, while controlling for alternative strategies.

On each trial participants chose to accept or reject the cumulative points on offer from sequences drawn from a unidirectional loop of 9 objects (Fig. 1B). Objects were defined by category (vehicle, fruit, animal) and colour (red, green, yellow). Each trial sequence depended on: a Start-object (where the sequence starts), a Reward-feature (a rule deciding which category or colour of objects are rewarded 2 points versus the default of -1 point) and a Stop-feature (a rule deciding which category or colour of object ends the sequence). Sequences with positive cumulative points should be accepted, those with negative points rejected. Points collected translated proportionally into monetary bonus (max £20).

**Fig 1:**
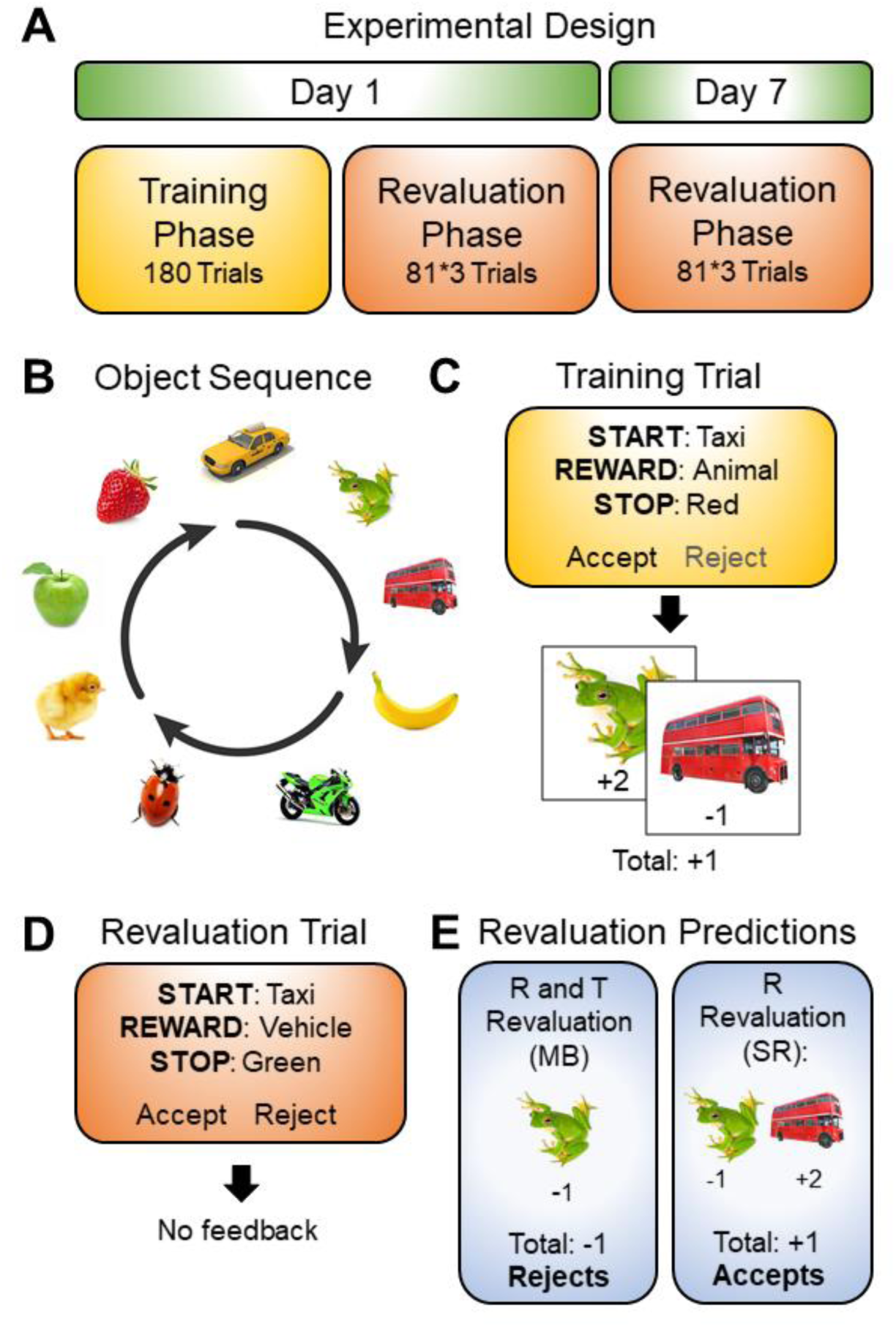
Experiment design. **A)** Training Phase on Day 1 is immediately followed by a Revaluation phase, which is repeated on Day 7. **B)** Nine objects, defined by 2 features (colour and category, e.g. frog: green animal), are organized in a unidirectional loop. **C-D)** Each trial’s sequence is determined by: 1. Start-object: where the sequence starts; 2. Stop feature: colour/category of the object which ends the sequence; 3. Reward feature: colour/category of objects that provide 2 points (others subtract 1 point). Participants choose ‘accept’ and take the total points (positive or negative) in the upcoming sequence (excluding the start object), or ‘reject’ and get zero points. (C) During Training each Start-object is paired with the same Stop/Reward Features and thus total reward. Each Start-object is presented 20 times in pseudorandom order. (D) Revaluation Trials pair each Start-object with every combination of Stop/Reward Features, creating 81 unique probes (72 new), with no feedback. **E)** Changes to Reward feature constitute reward (R) revaluation and changes to Stop feature constitute transition (T) revaluation. MB strategy is sensitive to R and T revaluation, whereas SR only to R revaluation.

During the Training phase (Fig. 1A), participants experienced 20 repeats of starting at each Start Object making an accept/reject choice based on the Start, Reward and Stop information, and then seeing the corresponding sequence of objects and points unfold sequentially (Fig. 1C). For each Start Object the Reward and Stop Features were unique and fixed, which meant that the same choice was always correct for the same Start object. All participants reached at least 85% accuracy in the final 20% of Training trials.

During Revaluation phases, which occurred immediately and repeated 7 days after Training (Fig. 1A), each Start-object was paired with every combination of Stop and Reward Features, creating 81 unique planning trials, of which 72 were new (Fig. 1D). Each unique trial type was repeated 3 times on each day. Participants made accept/reject choices and no further feedback was given. Relative to training, changes to Reward feature require reward (R) revaluation and changes to Stop feature require transition (T) revaluation. Both may impact which choice is correct (Fig. 1E).

### Identifying Sequential and non-Sequential Planners

We measured the use of each planning strategy in each participant by predicting the choices that MB, SR and “model-free” agents (MF i.e. habitual association of Start-objects with correct choice from Training) would make in the Revaluation-phase (Fig. 1E). These predictions assume that each strategy uses representations that were learned to asymptotic convergence in the training phase, which is supported by the high accuracy achieved by participants at the end of training. We use these predictions (+1: accept, -1: reject, 0: no prediction) to fit the Revaluation-phase choices of 30 participants performing the task in a magnetoencephalography (MEG) scanner, using logistic regression and Bayesian model comparison (see Fig. S1). The best fitting model, accounting for number of parameters, has three weighted factors: 1) general bias towards accepting; 2) MB prediction; 3) SR prediction. The MF predictor was not sufficiently explanatory to justify inclusion, and the SR prediction included the probability of each object following the given object during training, whether it was the Start-object or intermediate object in a sequence.

Regression weights were highly significant across the group for both the MB (t_29_=4.308, p<0.001) and SR predictors (t_29_=8.621, p<0.001; Fig. 2A, orange). We median split participants according to their MB weights, leaving two groups (each n=15): Sequential Planners (SPs; Fig. 2A, purple), who primarily rely on the MB (t_14_=6.500, p<0.001) but also an SR strategy (t_14_=5.923, p<0.001), and Non-Sequential Planners (nSPs; Fig. 2A, turquoise) who exclusively rely on the SR strategy (t_14_=6.461, p<0.001) and not the MB strategy (t_14_=0.653, p=0.262).

**Fig. 2:**
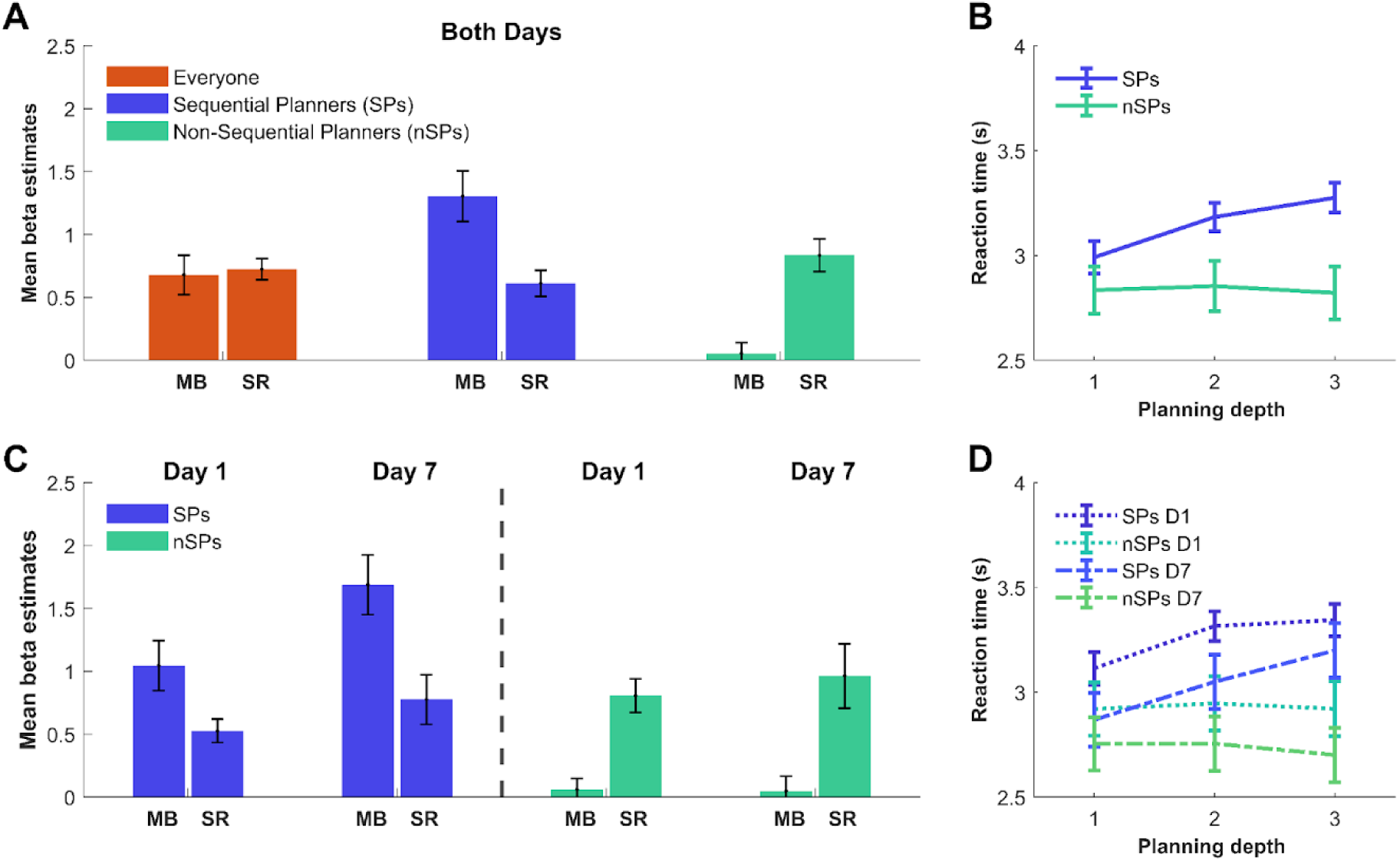
Planning behaviour. **A)** Mean parameter beta estimates for MB and SR regressors in the best fitting model for choices over both days and all participants (orange bars). Median splitting of participants based on their MB estimate produces 2 groups: Sequential Planners (SPs, purple bars) and non-Sequential Planners (nSPs, turquoise bars). **B)** Reaction time (s) as function of planning depth on each revaluation trial for SPs and nSPs. **C)** Mean MB and SR betas for SPs and nSPs, estimated for Day 1 and Day 7 separately. **D)** Reaction Time as function of planning depth on each revaluation trial, for SPs and nSPs, on Day 1 and Day 7. All error bars indicate SEs.

### Sequential Planners Take Longer to Plan Further

If sequential planning relies on step-by-step simulation, longer rollouts should require longer reaction times (RTs). This should not be true for non-sequential strategies. We therefore investigated the effect on RT of planning depth in the Revaluation-phase (i.e. 1, 2 or 3 objects, consistent with the Start and Stop objects on each trial). Consistent with the notion that only the SPs plan sequentially, there was a strong relationship in the SPs (F_1, 31.371_=19.418, p<0.001) but no significant relationship in the nSPs (F_1, 37.675_=0.0159, p=0.900) with a significant difference in RT modulation between the groups (F_1,29.509_= 21.382, p<0.0001) (Fig. 2B). RT modulation by planning depth that was almost perfectly correlated with their MB weight (r=0.87, p<0.001) across all participants. There was no significant effect on Revaluation phase RTs of the planning depth associated with each Start Object during the Training phase in either group (i.e. theoretical planning depth of the SR, see Supp. Table 1). These results indicate that only the strategies capable of transition revaluation (i.e. MB but not SR), which are seen in SPs, are sequential in nature.

### Sequential Planning Behaviour Improves with Consolidation

Being able to measure sequential planning, we next analysed the effect of consolidation upon this process by comparing behaviour on Day 1 v Day 7. Critically, there was no feedback during the Revaluation phases, and thus no learning. Day 7 compared to Day 1 only saw an increase in MB weights across everyone (t_29_=3.035, p=0.003). This increase primarily reflected MB weights increasing in SPs (t_14_=5.122, p<0.001) and significantly more so than in nSPs (t_14_=4.246, p<0.001; Fig. 2C). Consistent with these results we also saw effect of planning depth on RT across days in SPs (F_1,29.757_=5.4503, p=0.027) but not nSPs (F_1, 28.854_=0.478, p=0.494), and a significant difference between groups (F_1, 13905_=5.126, p=0.0235; Fig. 2D). We also found a general reduction in RT on Day 7 in both SPs (F_1,32.659_=22.154, p<0.001), and nSPs (F_1, 31.667_=5.635, p=0.0238). These results suggest that SPs make choices more consistent with MB planning over the course of consolidation (objectively and compared to nSPs), and that this reflects increasingly efficient step-by-step simulation or rollout through the task transition structure.

### MEG Reveals Consistent Planning Dynamics

Human neuroimaging aimed at measuring sequential activations have often relied on temporal cross-correlation or lagged regression of decoded state representations associated with early visual processing, via decoders trained 150-200ms after image onset (^26,28,28,29,33^). This approach makes assumptions about the mental representations of interest that may not be suited to step-by-step simulations likely reliant on more conceptual representations in the hippocampus or PFC ^34^. We therefore developed a different approach that uncovers sequential activity through a transition structure without making assumptions about the neural representations involved.

The approach is based on the notion that if planning entails sequential rollouts we should on average find consistent neural dynamics over the entire planning period across trials starting at the same object N (Fig. 3A). This is equivalent to the neural activity transitioning through the same series of distinct states based on where you start planning on the loop. Accordingly, we analyzed MEG data for the first 3s of planning in trials with the same start state N, that were randomly split into two halves, z-scored across trials (to represent differences between trials) and averaged within half. For each 0.1s time bin in each half resulting voxel vectors (1950 voxels in source space) were cross-correlated with every .1s time bin of the other half (Fig. 3A-C). Correlations were averaged over the N=1:9 and 50 random half-splits of trials (Fig. 3C). Consistent dynamics based on start state should look like a main diagonal (y-intercept=0) in the 2-D cross-correlation plot stretching throughout the 3s planning period (Fig. 3B), which is what we see in SPs (Fig. 4A) and nSPs (Fig. 4B). Similarity peaks at 0.3-0.4s with no difference between SPs and nSPs (t_14_=.432, p=.336). Mean similarity along the diagonal after this period (0.4 to 3 s) is marginally greater in SPs compared to nSPs (t_14_=1.310, p=.106) (Fig. 4C), hinting at more consistent neural dynamics in participants who plan sequentially.

**Fig. 3.**
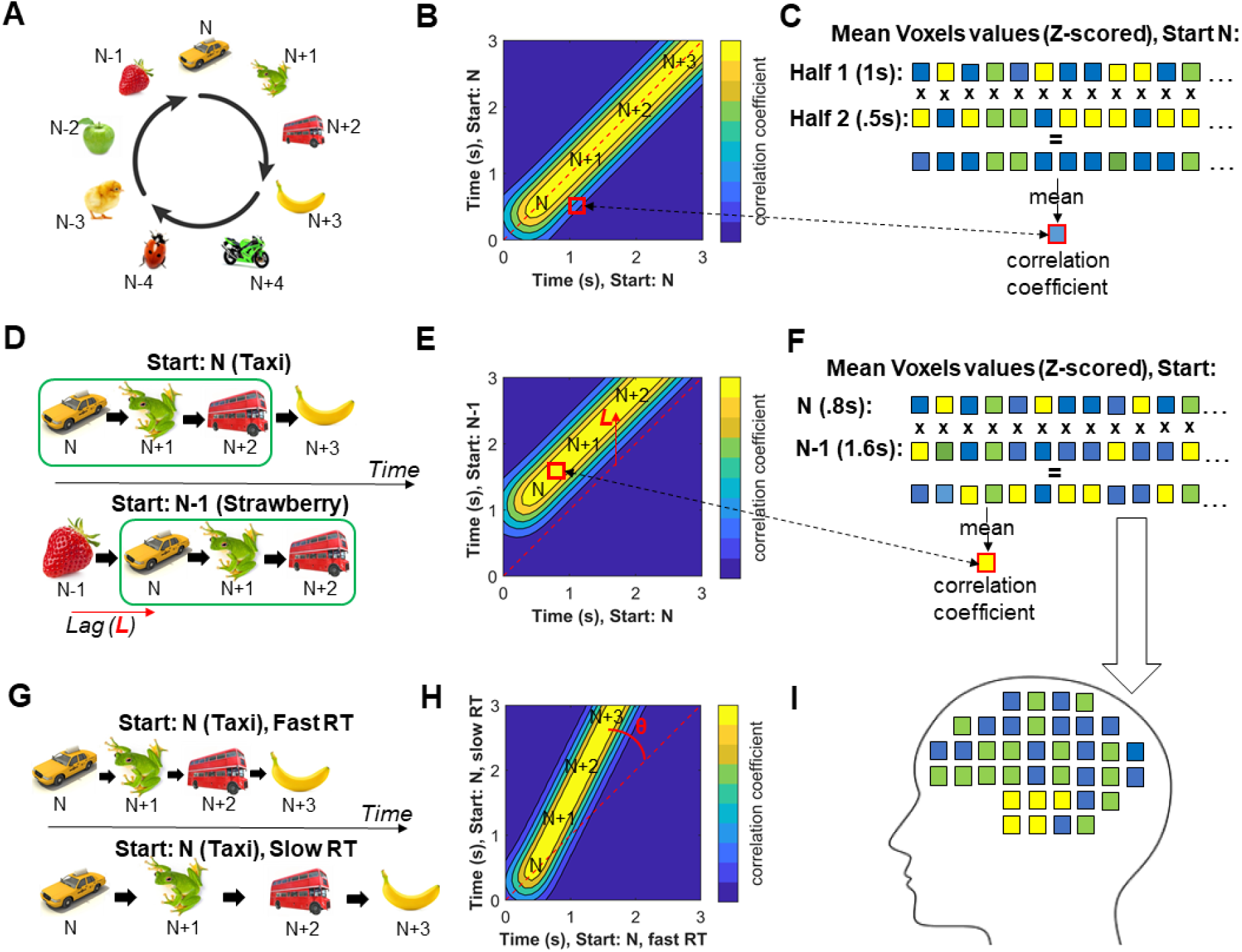
MEG Methods and Predictions for Neural Data: **A-C)** For each 0.1s in first 3s of Revaluation trials we correlate voxel values averaged over half of the trials (z-scored within half) with Start-object N (x-axis in B), with the corresponding values every 0.1s in the other half (y-axis in B) and average over the 9 Start-objects **(C)**. Consistent neural dynamics associated with sequential rollouts starting from N and progressing along the loop **(A)** should produce similarity along the zero-lag diagonal throughout the planning period **(B)**. **D)** Sequential rollouts through the loop transition structure should mean that neural patterns for Start-object N-1 should on average resemble those from Start-object N but shifted forward by a time-lag (L) (time to transition from one object to the next). Calculating the 2-D correlation plot for N relative to N-1 trials **(F)**, this should produce a diagonal with intercept L (**E**). **G)** If faster rollouts means faster planning, the rollouts should on average proceed faster when RT is shorter. Thus, comparing trials with slow and fast RT within participants (median split) when Start-object is N we should see a diagonal with a slope above 1, i.e. θ above zero **(H)**. **I)** To see which brain regions are driving the sequential rollouts (E) through the loop, we average all the voxel values in the upward shifted diagonal to find the voxels in the brain with larger values of the pair-wise products that form the correlation (F).

**Fig. 4.**
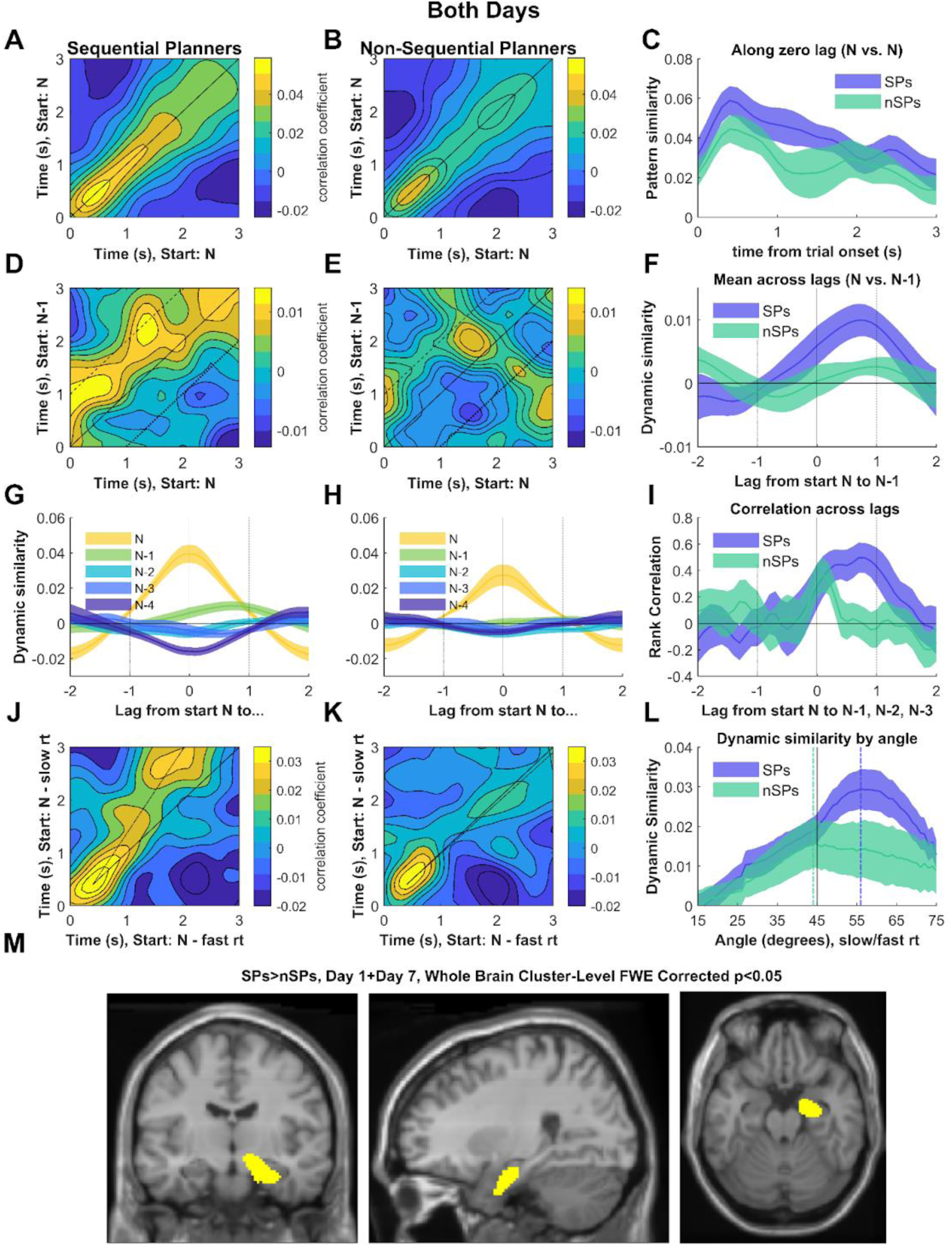
MEG Dynamics During Planning. **A-B)** Pattern similarity (group means) across voxels in SPs (A) and nSPs (B), comparing trials with the same Start-object (N), for all time-points in one half (x-axis) with the other half (y-axis) (cf. Fig. 3B) . **C)** Mean pattern similarity along zero-lag diagonal in SPs (A) and nSPs (B). Error bars indicate SE. **D-E)** Pattern similarity between trials with Start-object N (x-axis) and trials with Start-object N-1(y-axis) (cf. Fig. 3E) for SPs (D) and nSPs (E). **F)** Dynamic similarity (i.e. average pattern similarity along entire diagonal for each lag) between trials with neighboring Start-objects (N vs N-1, offset=1) in SPs and nSPs, **G-H)** Dynamic similarity across lags between trials with Start-objects with varied offsets (i.e. N vs N-1: distance=1, N vs N-2: distance=2, N vs N-3: distance=3) in SPs (G) and nSPs (H) I) Negative rank correlation between dynamic similarity and Start-objects offset in SPs (G) and nSPs (H) at each lag. **J-K)** Pattern similarity comparing trials with faster than median RT (x-axis) and slower than median RT (y-axis), all for Start-object N (cf. Fig. 3H). **L)** Dynamic similarity in SPs (J) and nSPs (K), along diagonals of varying angles through the plots J and K, passing through (0.3s, 0.3s). **M)** Source localization of the difference in dynamic similarity between SPs and nSPs for both days, averaged across 0.1-1s lags comparing trials with Start-object N to those with N-1.

### Sequential MEG Pattern Dynamics in Sequential Planners

To test if these consistent dynamics in MEG activity patterns truly reflected sequential rollouts through the loop, we repeated the above analysis, but now correlating trials with a given Start-object (N) with trials starting with the preceding object in the sequence (N-1). Sequential rollouts starting from object N-1 should on average progress through the same neural representations as rollouts starting from object N, delayed by some constant time-lag (L) (Fig. 3D-F). Consistent with such rollouts, in SPs only, the 2-D correlation plot shows high similarity on a diagonal that is shifted upwards by a constant time-lag for SPs (Fig. 4D). This indicates that trials starting with object N-1 instantiate the same ordered series of voxel activity patterns as trials starting with object N, but reach those patterns later. To quantify the time-lag we computed the similarity at each lag by averaging similarity along diagonals (slope=1) with different lags (y-intercepts). This “dynamic similarity” - the similarity averaged across the entire dynamics at a certain lag - shows significant lagged similarity for lags of 0.1 to 1 s, peaking around 0.8s (Fig. 4F), i.e. sequential neural dynamics advancing along the loop structure at a rate of one object roughly every 0.8s. This is not seen in nSPs (Fig. 4E), and the dynamic similarity over the range of lags 0.1 to 1s is significantly greater in SPs compared to nSPs (t_14_=2.513, p=0.011).

### Dynamic Similarity Between Planning from Start-objects with Varied Offsets

MEG dynamic similarity can be calculated for planning from Start-objects with a range of relative locations around the loop (comparing N to N, N-1, N-2, N-3 and N-4). In SPs (Fig. 4G), but not nSPs (Fig. 4H), similarity monotonically reflects distance between Start-objects on the loop. This relative similarity, in particular when comparing trials from Start-object N to those from N-1, N-2 and N-3 (for which planning involves overlapping sets of objects), is most prominent at lags of 0.1 to 1s. To quantify this we performed a rank correlation (since data have been z-scored across trials, only relative ranking is interpretable) between dynamic similarity and negative distance (1, 2 or 3) from N to N-1, N-2 and N-3, for lags 0.1-1s in each participant (Fig. 4I). We found a strong correlation for SPs (t_14_=4.600, p<0.001) which was significantly greater than nSPs (t_14_=2.128, p<0.026). Thus, dynamics which can distinguish between upcoming states based on distance from N, are sequentially following the dynamics during N by up to a second. Although no significant rank correlation was found for nSPs in that lag range (t_14_=0.837, p=0.208), there is a significant rank correlation in the 0-0.2s range (t_14_=2.795, p=0.007), which is significant 0s lag alone (t_14_=2.334, p=0.017), meaning that dynamics which can distinguish between upcoming states is simultaneous with dynamics during N. This suggests a mechanism for SR-based planning, whereby the neural representations of states that are experienced in succession become more similarly represented^35^ rather than reflecting sequential roll-outs.

### Planning Fast and Slow

Next, we investigated whether the speed of the MEG dynamic similarity reflected the speed of planning as judged by reaction times (RT). We split trials within participants, based on reaction time, controlling for day and planning depth, and plotted the similarity matrix for trials with the same Start-objects that had RTs below median versus those with RTs above median (Fig. 3G-H). These plots show the predicted steeper diagonal in SPs but not nSPs (Figs. 3J-K), indicating that speed of rollouts reflect planning speed. To quantify the slope we calculated the dynamic similarity along diagonals (going through 0.3s, 0.3s) across a range of slopes, finding the angle with peak similarity (Figure 3L). The slope of maximum dynamic similarity in SPs was 11°. For SPs (t_14_=3.386, p=0.002) but not nSPs (t_14_=0.103, p= 0.460) the mean angle θ (Fig. 3H) was significantly larger than 0, with a marginal difference between groups (t_14_=1.574, p=0.068).

### Planning to the End

Additionally, we wanted to verify that the sequential dynamics in SPs progressed all the way to the end of the sequence defined by the Stop feature of that trial. If that was the case we expected that the final state of the rollout should be decodable just before the response. Since we know that the speed of dynamics scales with RT, we normalized the neural data by resampling into 31 bins as proportions of the RT on that trial, and calculated similarity along the main 0-lag diagonal as a function of whether trials with the same Start object also shared Stop feature i.e. whether they have the same planning depth and therefore end on the same object. Thus, Same End similarity was calculated across pairs of trials with planning depths 1v1, 2v2 and 3v3, while Different End similarity was calculated across pairs of trials with planning depths 1v2, 2v3 and 1v3. Consistent with rollouts proceeding all the way to the end of the sequence, we found that in SPs, for the last 25% (last quartile) of the RT prior to response, pattern similarity was greater for trials that shared the same end state (t_14_=2.330, p=0.0176). This was not seen in nSPs (t_14_=0.498, p=0.313), with a near-significant difference between groups (t_28_=1.6455, p= 0.055). No such pattern was seen in the 1st, 2nd or 3rd quartile of normalized RTs in SPs (Fig. S2).

### Effect of Consolidation on Consistent Sequential Dynamics in Sequential Planners

Given the increases in MB estimates and RT modulation by planning depth over consolidation in SPs we wondered whether their neural dynamics more clearly reflected rollouts on Day 7 compared to Day 1. We found that the N vs N diagonal showed higher similarity throughout the first 0.4-3s of the trial (t_14_=1.863,p=0.043) for SPs (Fig. 5A-C), with a greater increase in SPs than nSPs (t_14_=1.761, p=0.050). In addition, the N vs N-1 dynamic similarity was significantly greater for lags in the range 0.1 to 1s on Day 7 compared to Day 1 for SPs (t_14_=1.813, p=0.046) (Fig. 5D-F) with a greater increase than for nSPs (t_14_=1.984, p= 0.033; Fig. 5F). We also saw a non-significant increase in rank correlation between mean dynamic similarity over 0.1 to 1s lags and the offsets between Start-objects on Day 7 compared to Day 1 (Fig. 5G-I; t_14_=0.784, p= 0.223). Lastly, the modulation of the speed of the dynamics by reaction time was enhanced on Day 7 compared to Day 1 in SPs (t_14_=2.187, p= 0.023) (Fig. 5J-L, Table S2), and significantly more than in the nSPs (t_14_=2.918, p= 0.006). The converging evidence indicates that rollouts became more stable, distinct and consistent with the task transition structure in SPs over the course of consolidation.

**Fig. 5.**
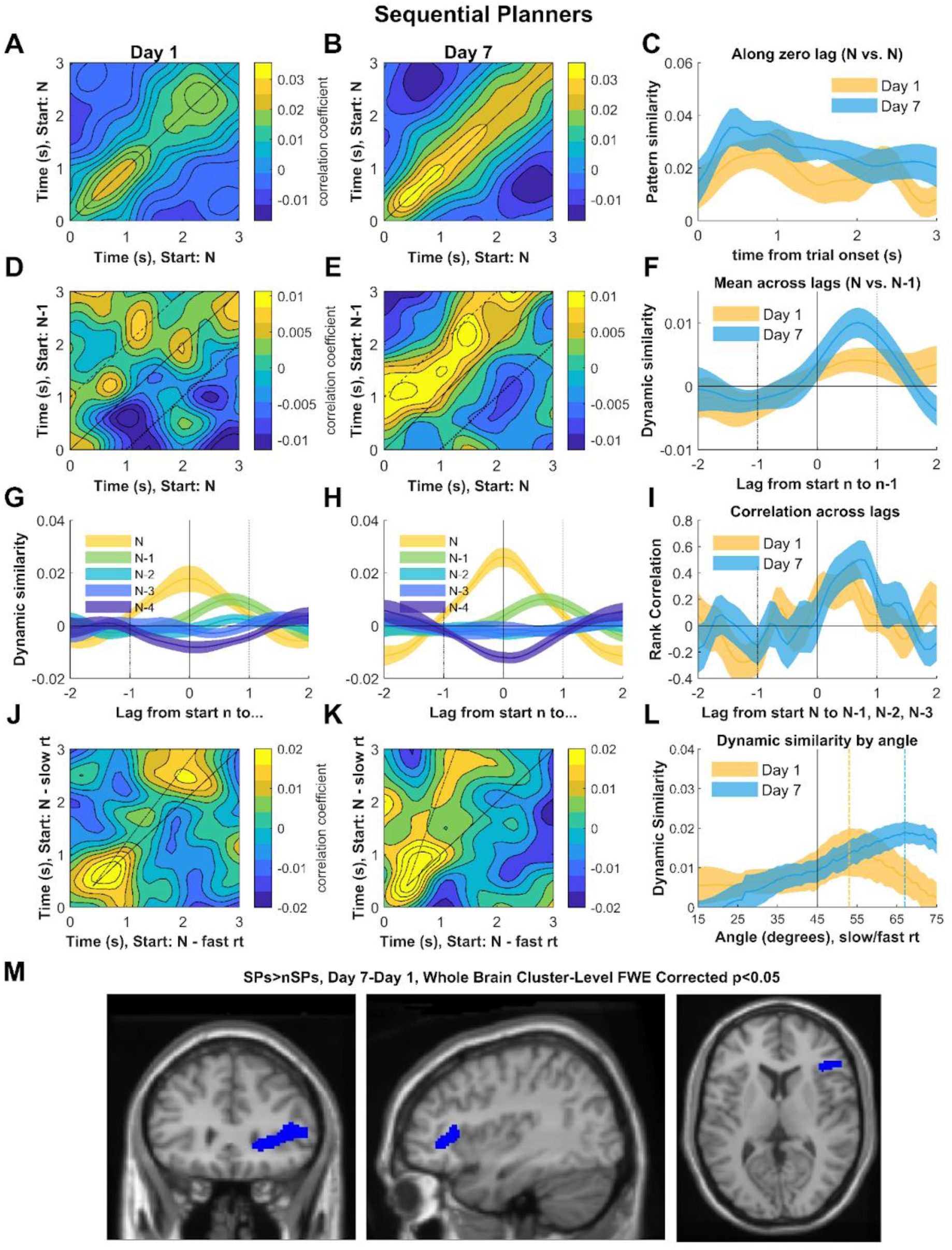
Consolidation of Sequential Planning in SPs. **A-L)** Same plots as Figure 4 but comparing Day 1 and Day 7 in SPs. (See Fig. S3 for nSPs) **M)** Source localization of the difference in dynamic similarity between SPs and nSPs for similarity on Day 7 minus Day 1, averaged across lags 0.1-1s comparing trials with Start-object N to those with N-1, A single cluster, overlapping the right prefrontal cortex was found.

### Source Localization of Sequential Planning Dynamics

We interrogated which brain regions are driving the lag-shifted dynamic similarity when comparing Start-objects N vs N-1, that was seen in SPs but not nSPs (Figs. S1E and 3D-E) and indicates sequential forward rollouts. To do this we looked for voxels contributing large pairwise products to the correlations on the diagonals corresponding to 0.1 to 1s lag (Fig. 3E-F & I). Once we had calculated this metric for each voxel in each participant over trials on both Day 1 and 7, we tested where values were higher in SPs relative to nSPs. Resulting t-statistics were then aligned to a standard brain, and the clusters of voxels surviving p<0.001 were tested at p<0.05 with family-wise error correction for the multiple comparisons using SPM . On^36^e cluster in the anterior temporal lobe overlapping the right anterior hippocampus and entorhinal cortex was found (Fig. 4M).

To reveal how neural substrates of sequential dynamics related to rollouts changed over the course of consolidation, we performed an identical analysis but instead tested where the dynamic similarity in SPs versus nSPs increased from Day 1 to Day 7. Only one cluster was found, in the right PFC, neighbouring OFC, lateral PFC and Broca’s area (Fig. 5M). No area saw significant decreases.

Further exploratory ROI analysis of the anterior temporal lobe cluster revealed that voxel-wise dynamic similarity did not change in SPs (t_14_=-0.021, p=0.843), or nSPs (t_14_=-0.901, p=0.3825) comparing Days 1 and 7. Further, looking in the right PFC ROI we found a significant increase in SPs (t_14_=2.985, p=0.009), but also, a significant decrease nSPs (t_14_=-3.320, p=0.005). These results support our hypotheses that in SPs, sequential rollouts take place the hippocampus and, following consolidation, increasingly the PFC.

## Discussion

Although flexible planning has most often been linked to prefrontal function^16,37^, recent theories ^4,16,38^ have emphasized hippocampal sequential rollouts as a possible planning mechanism, though limited direct evidence supports this proposal. Alternative accounts of hippocampal involvement in planning highlight either an indirect function of consolidation ^39–41^ or planning via non-sequential mechanisms ^31,42,43^. Here, we clarify the hippocampal contributions as online sequential rollouts at choice time, and propose a dynamic relationship between hippocampal and prefrontal mechanisms, linked through systems consolidation.

The time course of these rollouts, 100-1000ms per planning step, suggests slow and deliberate sequential planning. This contrasts with faster hippocampal processes like replay (40-200ms;^19,26,28,29,33^) which are more clearly associated with memory consolidation than online choice behaviour ^39,40^. Furthermore, the fact that pattern similarity when comparing trials with the same start state peaks 300-400ms after trial onset, as opposed to 150-200ms for image decoders ^26,28,29,33^, indicates that the rollouts likely involve conceptual rather than sensory representations^34^.

Critically, our findings reveal that consolidation strengthens both the behavioural markers of sequential planning and the neural signatures of rollouts, particularly in the PFC. These results contrast with a prevailing hypothesis in reinforcement learning which posits that offline consolidation processes serve to update either SR state predictions or MF action values, thereby reducing the reliance of flexible behaviour on sequential simulation at choice points^32,44^. Instead, our results show that consolidation increases behavioural flexibility at a choice time by strengthening the process of sequential future simulation through schemas of the transition structure. This is also consistent with behavioural evidence pointing to a role of consolidation in strengthening sequential representations^45,46^. Since hippocampal rollouts in SPs are undiminished a week later, our findings also indicate the hippocampus’s continued role in planning-related memory retrieval^47^. Nonetheless, the emerging PFC representations may alleviate the need for hippocampal processing, consistent with systems consolidation of flexible planning^22^.

Our results imply that consolidation transforms representations to afford flexible planning, although the nature of this transformation still requires clarification. MEG dynamics in MTL and PFC could potentially reflect sequential rollouts via a cognitive map or generative model initially learned in the hippocampus and entorhinal cortex ^6,48^ and consolidated into the prefrontal cortex for increased stability. Alternatively, the hippocampus could allow flexible sampling and recombination of individual episodic memories, to support both initial planning ^49–51^ and the eventual consolidation of a more abstract generative model in prefrontal cortex, for improved flexibility and generalizability ^8–15,52^. This latter possibility aligns with the distinction between the use of specific associative information stored “in-weights” (in hippocampus) and “in-context” (in PFC) processing determined by initial activity patterns playing out through a network architecture that captures the general structure of the task ^52–54^. Future work will be required to distinguish these possibilities and whether the consolidation demonstrated here is driven by reactivation during task performance or rest ^13,28,40,41,55^.

A principal theme of cognitive neuroscience has concerned the distinction between flexible goal-directed planning and automatic habits ^16,23,24,37^. Here we explore a new dichotomy within flexible strategies that contrasts sequential planning, which relies on step-by-step simulation using a transition structure, against non-sequential planning which associates states based on occurrence statistics without encoding sequential order^30–32^. While the neural data from sequential planners clearly demonstrate an algorithmic process for MB evaluation involving rollouts, we speculate that SR evaluation is accomplished by associative mechanisms^35^, which cause states presented together to take on overlapping representations (see Fig. 4I for nSPs around lag 0). The significant variance in these planning strategies across participants might reflect both incidental factors^56^ and more permanent biases in learning and memory mechanisms or traits that affect capacity for sequential planning ^57–59^. It will be of great importance in future work to determine how flexibility is impacted by consolidation, as only participants who initially developed representations for rollouts, were capable of later consolidating these for improved planning capabilities. Furthermore, understanding these distinctions may be critical for making sense of the development of mental health disorders, like OCD, linked to dysregulation of flexible decision-making ^60^.

## Acknowledgements

Thank you to Daniel Bush, George O’Neill, Oded Bein and Gareth Barnes for insightful comments on the manuscript. We are also thankful to Jesse Geerts and Rachel Bedder for helpful conversations, and the imaging support staff at the Wellcome Centre for Human Neuroimaging. ChatGPT 4o was used for language processing of the manuscript.

## Ethics

Ethical approval was approved by the local research ethics committee at University College London. All participants gave written informed consent and were compensated for their participation.

## Author contributions

Conceptualization: OMV, EMR, NB

Methodology: OMV, EMR, NB

Investigation: OMV

Visualization: OMV

Funding acquisition: NB

Project administration: OMV

Writing – original draft: OMV

Writing – review & editing: NB, EMR, OMV

## Supplementary Methods

### Task details

The experiment is split in two phases: Training and Revaluation. The Training Phase on Day 1 is immediately followed by a Revaluation phase, which is repeated on Day 7. The experimental design is based on an object sequence of nine objects organized in a sequential unidirectional loop which defines the state space and transition structure of the task. The object sequence is pseudo randomized and fixed for each participant. Each object is uniquely defined by 2 features: category (fruit, vehicle, animal) and colour (red, yellow, green), e.g. frog (green animal).

On each trial, 3 pieces of information deterministically decides the upcoming sequence: 1. *Start Objec*t: where sequence starts; 2. *Stop Feature*: colour/category of object which ends sequence. The sequences can be of length 1,2 or 3 objects, not including start object; 3.

*Reward Feature*: objects from this category/colour provide 2 points (all other objects deduct 1 point). Points are cumulative across the sequence on that trial, excluding the start state. For each participant, one feature always represents either *Stop* Or *Reward*, counterbalanced across participants. Each trial, participants either choose ‘accept’ and take the cumulative points (positive or negative) on the upcoming object sequence, or ‘reject’ and get zero points. Participants are told that they will receive a monetary bonus that stands in direct proportion to the number of points won (max £20).

During the training phase, each *Start Object* is always paired with the same Stop/Reward Features, i.e. produces the same sequence and total reward. Also, during the training phase, following accept/reject choice the Start Object-specific sequence will unfold regardless of choice, providing feedback on every trial. Participants have up to 10s RT to make a choice during the Training phase. If no choice is made in 10 seconds, the trial times out and there is no feedback for that trial. If a response was made and RT was above 3 seconds, the sequence played out immediately after choice. If RT was below 3s, the screen remained unchanged until 3s had passed since the beginning of the trial, after which the sequence unfolded. Trials with each *Start Object* are presented 20 times (i.e. 180 trials during training Phase), in pseudo-random order. During Training sequences were arranged such that 4/9 Start objects lead to a sum of >0 points (should accept), 4/9 lead to a sum of <0 points (should reject) and 1/9 lead to a sum of 0 points (should accept or reject). The point total is never presented to participants but needs to be added up sequentially over the trial.

During the Revaluation Phase each Start object is subsequently paired with every combination of Stop/Reward Features, creating 81 unique planning probes, of which 72 are new (Fig. 1C). On both Day 1 and 7, each of the 81 probes are repeated in 3 trials. No feedback (sequential objects or rewards) follows choices during the revaluation phase. For this phase, max rt is 5s, with a 3s minimum for the trial to end. Participants are told that even though they wouldn’t receive feedback they will still be rewarded in proportion to the points won, and that on Day 7 the potential for monetary bonus in the Revaluation Phase is identical on Day 1.

Prior to experiment beginning, participants were given an instruction tutorial and 10 question quiz. Any incorrect quiz answers requires re-examining the tutorial until all questions are answered correctly. During the tutorial participants were briefly exposed to the full object sequence initially by experiencing the loop 3 times sequentially, for each object making a binary choice between the correct object and one of the other objects. Participants also performed practice trials, first being exposed to only the object sequences without rewards and then for each Start object making choices with feedback. The example trials are identical to the trials from the Training phase, with the same fixed Reward and Transition features with each Start object. All participants are also informed that within the 9 object sequence, the feature which represents Stop for that participant (e.g. colour) will be repeated in the same sequence (e.g. red, yellow, green, red, yellow, green, red, yellow, green). The reward feature does not have a predictable pattern.

When participants returned on Day 7 we confirmed that they could complete the quiz without errors but they were not reminded or instructed on the object sequence nor saw any pictures of the objects. We also ran a localizer task where each object image was presented 45 times each for .7s in pseudorandom order prior to the tutorial and at the end the the Revaluation Phase on Day 7. Additionally, before and after each Revaluation phase we included a 3min rest period where participants relaxed while a fixation cross was on screen.

### Logistic Regression Analysis Choice Data

We measured the use of each planning strategy in 30 participants performing the task in a magnetoencephalography (MEG) scanner, by predicting the choices that MB (R and T revaluation), SR (R revaluation) and MF (no revaluation) agents would make in the Revaluation-phase based on what they learning during Training phase and novel Reward/Stop features. These predictions assume that each strategy uses representations that were learned to asymptotic convergence in the training phase, which is supported by the high accuracy achieved by participants at the end of training. We use these predictions (+1: accept, -1: reject, 0: no prediction) to fit the Revaluation-phase choices using logistic regressions fit separately to each participant’s data, implemented using fitglme in MATLAB. If, for insurance, based on the correct transition structure and the specific Start object, Stop and Reward features of a trial, a model based agent expects -3 points, the MB prediction for that trial is -1 (reject).

### Model Comparison

Model fits are evaluated using Bayesian Information Criterion (BIC) (Stone, 1979) that measures the trade-off between model fit and complexity of the model. The best fitting model has three weighted factors: 1) general bias towards accepting; 2) MB prediction; 3) SR prediction (Fig. 3). The MF predictor was not sufficiently explanatory to justify inclusion. The SR prediction meant the probability of every object following a given object during training, whether the given object was the Start-object or intermediate object in a sequence. Additional models were tested with only the SR for each object when that object was the Start object (encountered as a word) (SRstart), or models with both SRs and another SR where the object was an intermediate object in another sequence, encountered as an image (SRint). Significance of group effects were tasted with one-tailed t-tests (>0 for individual groups, MB>SR, for group comparisons, Day 7 > Day 1 for consolidation tests). Participants were separated into groups based on median split of MB beta estimates from individual logistic regression.

### Linear Regression Analysis of RT Data

We also investigated the effect on RT of planning depth in the Revaluation-phase (i.e. 1, 2 or 3 objects, consistent with the Start and Stop objects on each trial). The RT effect was tested with a mixed linear regression with participant ID as random factor implemented with fitglme in MATLAB (standard errors computed using the Satterthwaite approximation to the degrees of freedom). In addition to planning depth and covariates, like group (SPs vs nSPs) or Day, we also included a variable indicating whether participants accepted or rejected, which significantly explained RT. We first tested planning depth (Dreval) implied post T revaluation (MB planning depth), i.e. based on the Start object and Stop feature on each Revaluation trial (Fig. S1). We Then tested planning depth (Dtrain) during training (SR planning depth), i.e. based on the Stop feature that was linked to each Start object during the Training Phase.

### MEG Preprocessing

MEG data collection and pre-processing MEG data were acquired using a 275-channel axial gradiometer system (CTF Omega, VSM MedTech) at a sample rate of 600 Hz. During the recording, head position coils (attached to nasion and left and right pre-auricular sites) were used for anatomical co-registration, and eye tracking was performed using an Eyelink 1000 system (SR Research). Raw MEG data were imported into SPM1244 and downsampled to 100 Hz before an independent component analysis (ICA) was implemented in FieldTrip45 and EEGLAB and any components with a correlation to eye-movement estimates were higher than .2 Finally, a fifth-order, zero-phase Butterworth filter was used to remove slow drift (1 Hz high-pass) and mains noise (48–52 Hz notch) from the recordings. Our analyses focused on planning periods from the trial onset until response time after max 5 s. Planning ‘epochs’ were defined as [-2 5] s windows around each trial onset. Once the MEG data had been epoched, artifact trials and channels were automatically identified and removed using an underlying outlier test (with a threshold of α=0.05). Trials with no response (RT ∼<5s) were also removed. Source localization of the full power spectrum during the Revaluations phase planning epochs was finally performed in SPM12 using the Linearly Constrained Minimum Variance beamformer from the DAiSS toolbox, with a single-shell forward model and sources evenly distributed on a 10 mm grid co-registered to Montreal Neurological Institute (MNI) coordinates. This resulted in a set of linear weights for each participant that could generate a time series from sensor-level data into 1950 voxels in source space during each planning epoch.

### MEG Analysis

If planning entails sequential rollouts we should see consistent neural dynamics over the planning period across trials starting at the same object N in the loop (Fig. 3A). Accordingly, we analyzed MEG data from the first 3s of the planning period of each trial of the Revaluation phases, using a variation on cross-temporal RSA (Kriegeskorte and Kievit, 2013, Luyckx et al 2019) (Fig. 3A-C). After preprocessing, the data for each trial comprised 30 MEG voxel vectors, one per 0.1s timepoint. We divided the trials randomly into two halves. For each half, for each timepoint, we z-scored the MEG voxel values across trials (to represent differences between trials rather than absolute values) and averaged the vectors corresponding to the same start Start-Object N. The voxel vector for each timepoint in each half were then correlated (mean of pair-multiplied z-scored vectors), with each timepoint’s corresponding vectors with the same Start-object N from the other half, comparing all timepoints in one half with those all timepoints in the other (results are averaged over the N=1:9 and 50 random half-splits of trials) (Fig. 3C). Consistent dynamics should look like a diagonal in the 2-D correlation plot stretching throughout the planning period (Fig. 3B).

To test if these consistent dynamics in MEG activity patterns truly reflected sequential rollouts through the loop, we repeated the above analysis, but now correlating trials with a given Start-object (N) with trials starting with the preceding object in the sequence (N-1), for all timepoints and averaging across Start-objects. Sequential rollouts starting from object N-1 should progress through the same neural representations as rollouts starting from object N, delayed by some constant time-lag (L) (Fig. 3D-F) which should reflect in the 2-D correlation plot as a diagonal that is shifted upwards. The size of the upward shift should indicate the time lag it takes to go from one state to the next during rollouts. To quantify the time-lag we computed the “dynamic similarity” at each lag by averaging similarity along diagonals (slope=1) i.e. the entire dynamics, for different lags (y-intercepts).

MEG dynamic similarity can be calculated for planning from Start-objects with a range of relative locations around the loop (comparing N to N, N-1, N-2, N-3 and N-4). In SPs (Fig. 3G), but not nSPs (Fig. 3H), similarity monotonically reflects distance between Start-objects on the loop. This relative similarity, when comparing trials from Start-object N to those from N-1, N-2 and N-3 ( for which planning involves overlapping sets of objects), can be quantified with rank correlations (since data have been z-scored across trials, only relative ranking is interpretable) between dynamic similarity and negative distance (1, 2 or 3) from N to N-1, N-2 and N-3, for different lags in each participant (Fig. 3I).

Next, we investigated whether the speed of the MEG dynamic similarity reflected the speed of planning as judged by reaction times (RT). We split trials within participants, based on reaction time, controlling for Day and planning depth (equally sampling trials from both days and all planning depths in each half), and plotted the similarity matrix for trials with the same Start-objects that had RTs below median versus those with RTs above median (Fig. 3G-H). A faster dynamic for fast RT trials should here be reflected as a slope above 1. To quantify the slope we calculated the dynamic similarity along diagonals (going through 0.3s, 0.3s) across a range of slopes. To test statistically if the slope is significantly higher than 1 (angle>0) we found the angle of maximum dynamic similarity for each participant after having 2D smoothed the similarity plot with 1s gaussian Kernel, and performed a one-tailed t tests to see if the angle was greater than zero across groups.

Additionally, we wanted to verify that the sequential dynamics in SPs progressed all the way to the end of the sequence. If that was the case we expected that the final state of the rollout should be decodable just before the response in SPs. Since we knew that the speed of dynamics scaled with RT, we therefore normalized the neural data by resampling the data into 31 bins as proportion of the RT on that trial, and calculated similarity along the main 0-lag diagonal as a function of whether trials with the same Start object also shared Stop feature i.e. whether they have the same planning depth and therefore end on the same object. Thus, Same End similarity was calculated as the average of planning depth 1v1, 2v2 and 3v3 trials, and Different End similarity was calculated as the average of planning depth 1v2, 2v3 and 1v3 trials and we tested if similarity was greater for same end trials in the last quartile of RT using a one-tailed t test.

Unless specified all statistics were performed on unsmoothed data. Figures are smoothed in 2D for illustrative purposes with a 1s gaussian Kernel.

### Source Localization

We interrogated which brain regions are driving the lag-shifted dynamic similarity when comparing Start-objects N vs N-1, that was seen in SPs but not nSPs (Fig. 3E and 3D-E) and indicates sequential forward rollouts. To do this we looked for voxels contributing large pairwise products to the correlations on the diagonals corresponding to 0.1 to 1s lag (Fig. 3E-F & I). Once we had calculated this metric for each voxel in each participant over trials on both Day 1 and 7, we tested where values were higher in SPs relative to nSPs, respectively. Resulting t-stats were then aligned to a standard brain, and the size clusters of voxels surviving p<0.001 and p<0.05 with family-wise error correction for the multiple comparisons using SPM.

We next performed an exploratory ROI analysis of how neural signatures of sequential dynamics changed from Day 1 to 7. This was done by extracting mean N vs N-1 dynamic similarity, the range 0.1 to 1s lag across voxels in the clusters in hippocampus and PFC separately for each day.

## Supplementary Results

**Fig. S1:**
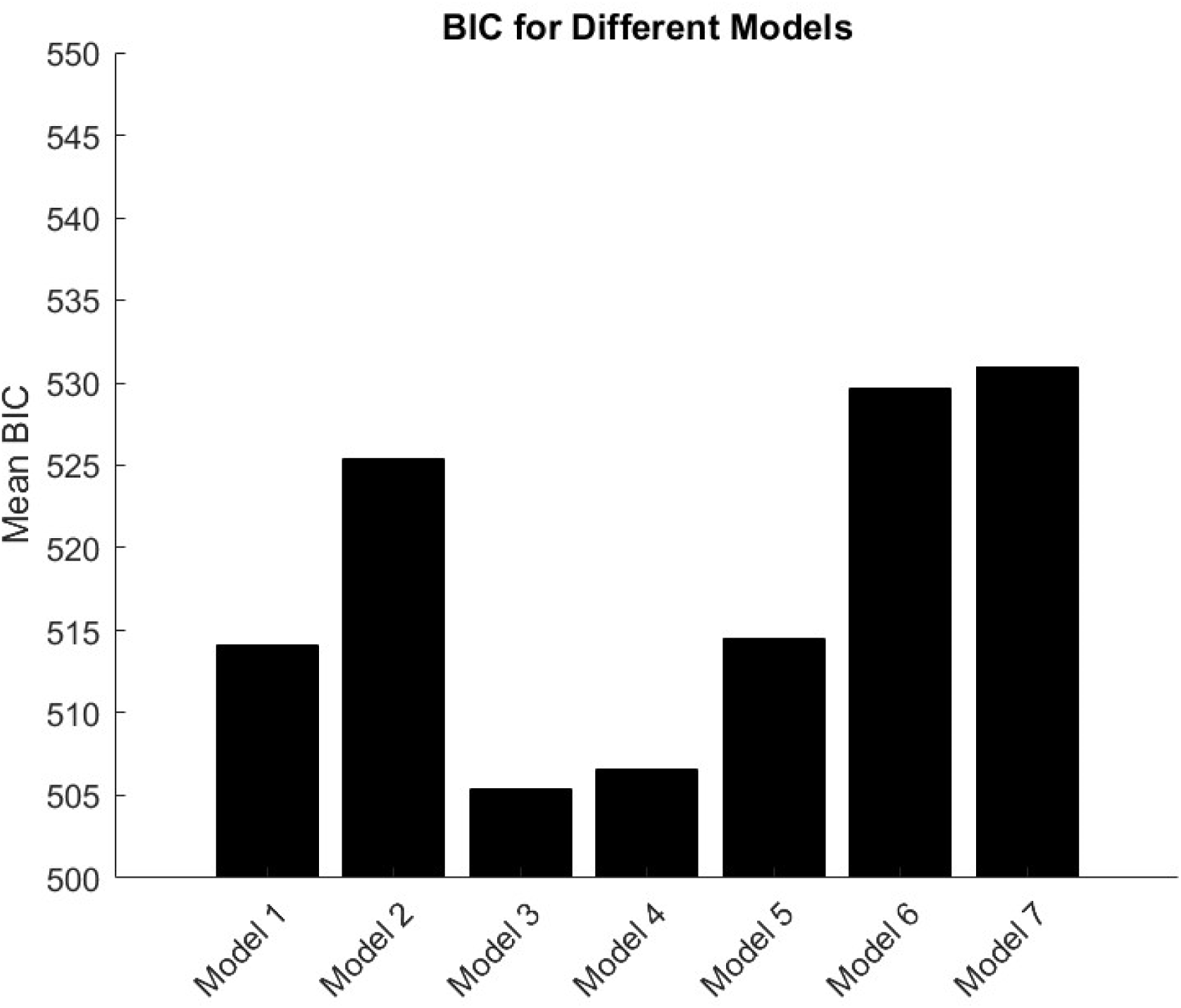
Mean BIC across participants. Fitted on Revaluation data for Day 1 and 7. For choice prediction regressors expected reward>0 coded as 1, expected reward<0 coded as -1 and expected reward=0 coded as 0. Tested models: **Model1**: 1. Accept bias; 2. MB **Model2:** 1. Accept bias; 2. SR **Model3:** 1. Accept bias; 2. MB; 3. SR* **Model4:** 1. Accept bias; 2. MB; 3. SR; 4. MF **Model5**: 1. Accept bias; 2. MB; 3. SRstart **Model6:** 1. Accept bias; 2. MB; 3. SRstart; 4. Srint **Model7:** 1. Accept bias; 2. MB; 3. SRstart; 4. SRint; 5. MF *SR denotes choice prediction calculated with successor states for every state when that state is the Start state (SRstart) or intermediate state in another sequence (SRint).

**Fig. S2:**
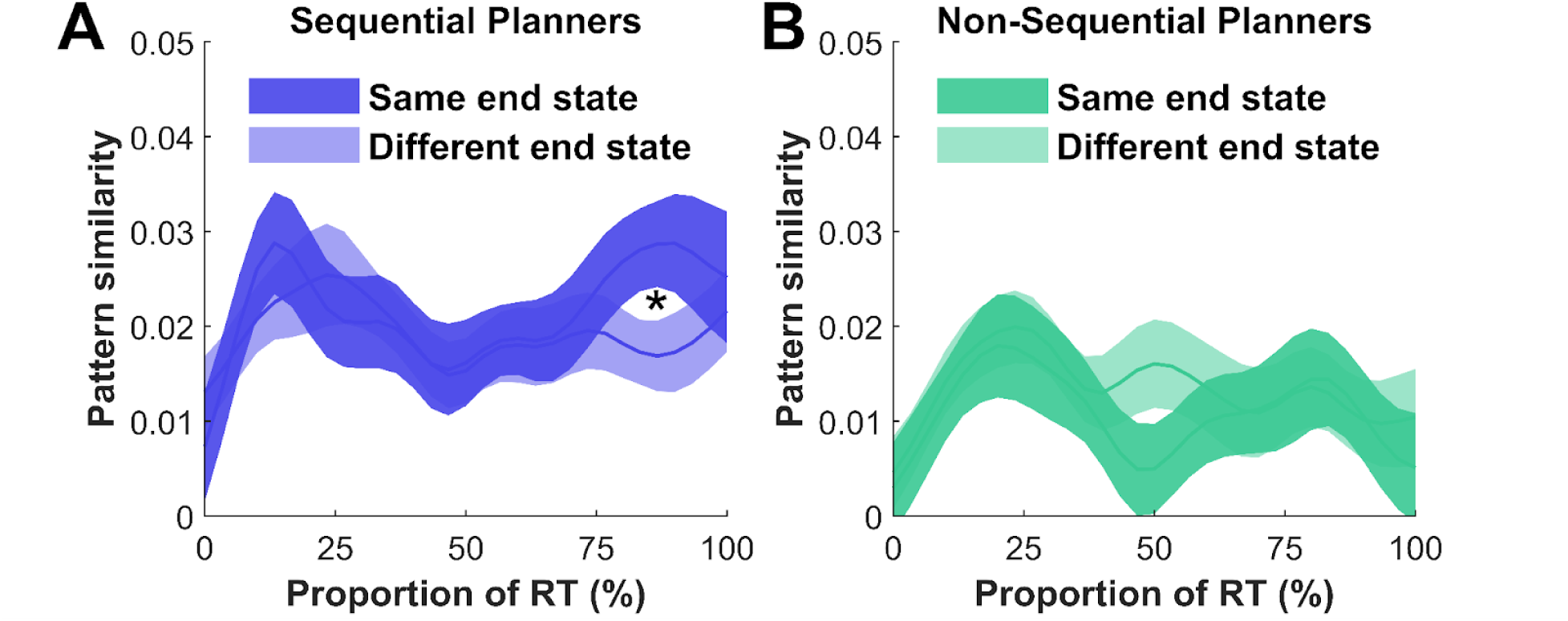
Separating Dynamics Based on End State. Fitted on Revaluation data for Day 1 and 7. Pattern similarity along 0-lag showing mean (and SE) pattern similarity in SPs (A) and nSPs (B) for neural data normalized as proportion of the RT on that trial. Compared trials share Start objects but are additionally separated based on whether they have a same or different end state.

**Fig. S3:**
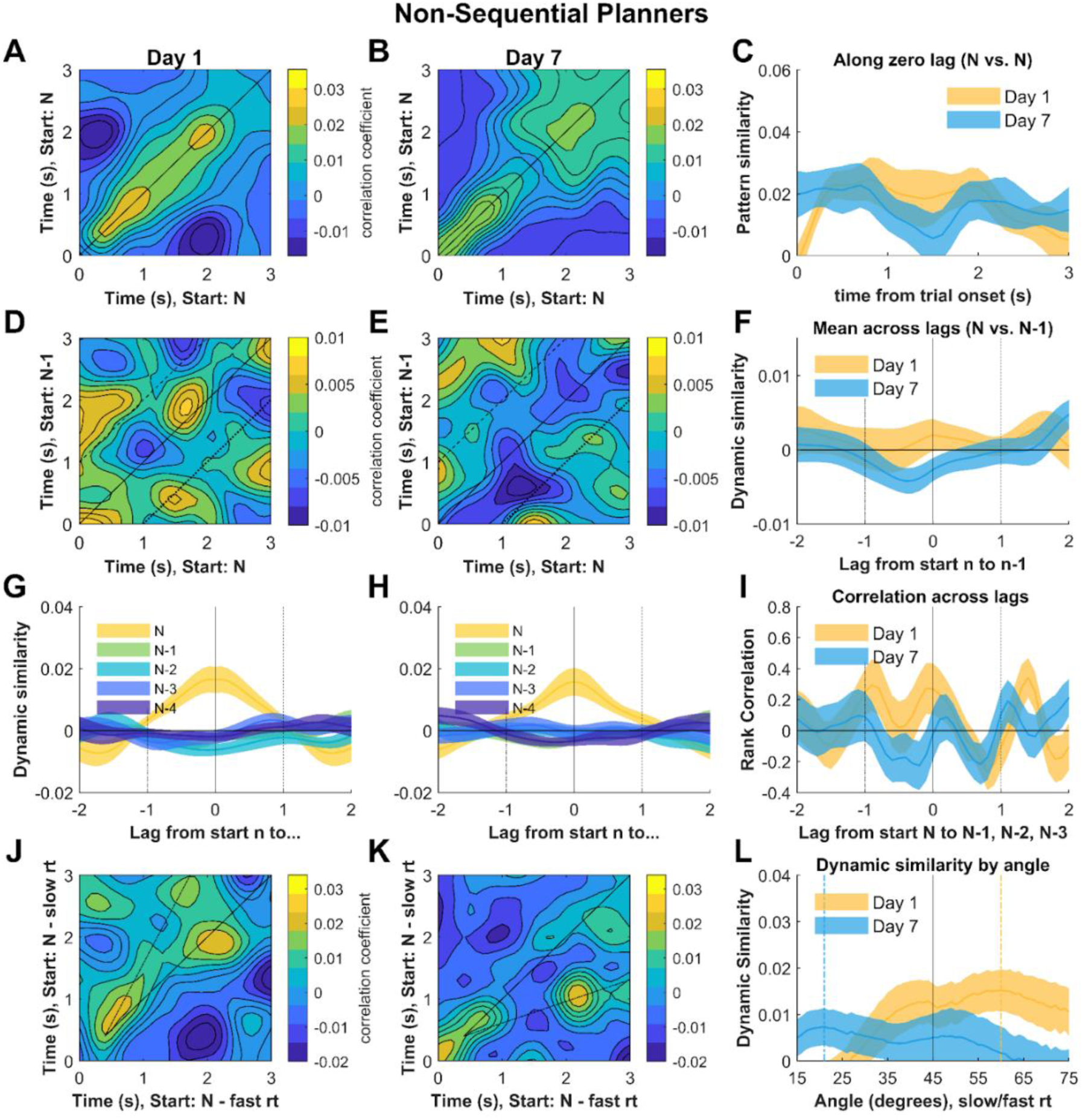
Consolidation of Sequential Planning in nSPs. **A-L)** Same plots as Figure 4 A-L but comparing Day 1 and Day 7 in nS

**Table S1:** Effect of revaluation (Dreval) and training (Dtrain) planning depth on RT, tested with mixed linear regression of (Satterwaite approximation for correct degrees of freedom) in SPs and nSPs. Dtrain reflect mean of depths of SRstart and SRint. Trial_accept is a variable coded 0 and 1 based on whether participants accept or reject.

| Name | Estimate | SE | Fstat | DF1 | DF2 | pValue |
| --- | --- | --- | --- | --- | --- | --- |
| Trial_accept | -103.81 | 36.273 | 8.19 | 1 | 30.202 | 0.007 |
| SPs | 2891.1 | 113.52 | 648.57 | 1 | 31.452 | 1.4687e-22 |
| nSPs | 2854.5 | 113.53 | 632.22 | 1 | 31.501 | 2.0452e-22 |
| Dreval:SPs | 145.59 | 23.913 | 37.065 | 1 | 30.918 | 9.654e-07 |
| Dtrain:SPs | 32.033 | 51.666 | 0.3844 | 1 | 31.638 | 0.53969 |
| Dreval:nSPs | -7.5806 | 23.895 | 0.10064 | 1 | 30.789 | 0.7532 |
| Dtrain:nSPs | 74.558 | 51.376 | 2.106 | 1 | 31.012 | 0.15676 |

**Table S2:** Effect of revaluation (Dreval) and training (Dtrain) planning depth and Day (1vs7) on RT, tested with mixed linear regression of (Satterwaite approximation for correct degrees of freedom) in SPs and nSPs. Dtrain reflect mean of depths of SRstart and SRint. Day coded as 1 for Day 7, otherwise zero.Trial_accept is a variable coded 0 and 1 based on whether participants accept or reject.

| Name | Estimate | SE | Fstat | DF1 | DF2 | pValue |
| --- | --- | --- | --- | --- | --- | --- |
| Trial_accept | -104.93 | 35.694 | 8.642 | 1 | 30.236 | 0.0062 |
| SPs | 119.37 | 25.661 | 658.49 | 1 | 31.112 | 1.6904e-22 |
| nSPs | 119.38 | 24.618 | 606.02 | 1 | 31.167 | 5.4794e-22 |
| Dreval:SPs | 117.51 | 4.406 | 19.418 | 1 | 31.371 | 1.1427e-4 |
| Dtrain:SPs | 8.483 | 67.139 | 0.0159 | 1 | 37.675 | 0.900 |
| Dreval:nSPs | -0.229 | 26.637 | 7.42e-05 | 1 | 31.219 | 0.993 |
| Dtrain:nSPs | 72.612 | 66.87 | 1.1791 | 1 | 37.125 | 0.285 |
| SPs:Day | -337.16 | 71.632 | 22.154 | 1 | 32.169 | 4.5992e-05 |
| nSPs:Day | -169.35 | 71.337 | 5.635 | 1 | 31.667 | 0.023837 |
| Dreval:SPs:Day | 56.47 | 24.188 | 5.4503 | 1 | 29.757 | 0.026507 |
| Dtrain:SPs:Day | 40.125 | 81.466 | 0.242 | 1 | 194.83 | 0.6229 |
| Dreval:nSPs:Day | -16.587 | 23.981 | 0.478 | 1 | 28.854 | 0.49465 |
| Dtrain:nSPs:Day | 6.957 | 80.993 | 0.007 | 1 | 188.77 | 0.93164 |

